# The Mutated p.H222P A-type Lamins Drive Loxl2-Mediated Extracellular Matrix Remodeling in Both Patient-Derived Cardiomyocytes and Mouse Models of Dilated Cardiomyopathy

**DOI:** 10.1101/2025.01.10.632312

**Authors:** Marie Kervella, Cecile Peccate, Zoheir Guesmia, Fiorella Grandi, Nathalie Mougenot, Anne Forand, Azzouz Charrabi, Guy Brochier, Ramaroson Andriantsitohaina, Albano C. Meli, Antoine Muchir

## Abstract

*LMNA* cardiomyopathy, caused by mutations in the *LMNA* gene, is a severe form of dilated cardiomyopathy characterized by arrhythmias, contractile dysfunction, and increased myocardial fibrosis, which impairs left ventricular function and predisposes to heart failure. While the disease has been well characterized, a lack of insight into the pathogenesis impeded the development of therapies. We here used patient-derived *LMNA* p.H222P cardiomyocytes (hiPSC-CMs) and their isogenic controls and a *Lmna*^H222P/H222P^ mouse model to dissect abnormal cardiac mechanisms leading to the development of the disease. We showed that *LMNA* p.H222P hiPSC-CMs exhibit elevated diastolic calcium levels and hypocontractility. They displayed nuclear shape abnormalities, a hallmark of *LMNA* cardiomyopathy, associated with altered chromosome spatial organization and gene expression profiles. Using transcriptomic analysis, we further revealed that genes related to cardiac extracellular matrix (ECM) remodeling, deposition, and components are dysregulated in both *LMNA* p.H222P hiPSC-CMs and mutated mice, suggesting a conserved pathogenic mechanism across species. Conversely, molecular inhibition of Loxl2, a key component of the ECM establishment, preserved the cardiac function *in vivo*. Taken together, our findings suggest that targeting Loxl2 could be a promising therapeutic strategy to maintain cardiac function in *LMNA* cardiomyopathy.

## INTRODUCTION

Inherited cardiomyopathies are a major cause of heart disease, often with an onset in adolescence or early adult life. Among these diseases, dilated cardiomyopathies (DCM) are characterized by left ventricular dilatation, systolic dysfunction, myocyte death, and myocardial fibrosis^1^. More than 40 disease genes have been identified; the most common mode of inheritance is autosomal dominant transmission, although autosomal recessive and X-linked forms have been described^2^. Mutations in *LMNA*, the gene encoding the nuclear A-type lamins, a key component of the nuclear lamina, are responsible for DCM (i.e. *LMNA* cardiomyopathy)^3^. These mutations represent the second most common genetic cause of familial DCM, making their pathogenesis a critical study area. *LMNA* cardiomyopathy exhibits a rapid clinical progression, characterized by a higher incidence of aggressive arrhythmias and quicker development toward heart failure compared with the majority of other cardiomyopathies^4^, as well as cardiac fibrosis^5–7^.

The nuclear lamina is a dense network of filamentous proteins that resides on the inner nuclear membrane, providing structural support to the nucleus and influencing the genome organization within the cell^8^. While the genetic causes have been identified, the molecular and cellular mechanisms behind *LMNA* cardiomyopathy remain poorly described^9^, hindering the development of treatment. Despite the widespread expression of A-type lamins in various tissues, mutations affect soft and stiff tissues. This paradox has led researchers to propose that changes in nuclear rigidity may influence gene expression, chromatin organization, and signal transduction pathways^10^. Mutations in *LMNA* result in nuclear abnormalities that disrupt the structural integrity of the nuclear envelope, a hallmark of *LMNA* cardiomyopathy^11,12^. These alterations in nuclear morphology have implications for cellular function and are thought to underlie the tissue-specific pathologies observed in the disease, though the precise mechanisms remain poorly understood. Understanding how *LMNA* mutations result in cardiac-specific phenotypes could provide insight into disease progression and cellular dysfunction underlying mechanisms.

The nuclear lamina is not only structurally important, but it also interacts with specific regions of genomic DNA called lamina-associated domains (LADs), which are chromatin regions that are physically tethered to the inner nuclear membrane^13,14^. LADs are typically transcriptionally repressed, and their positioning within the nucleus plays a significant role in regulating gene expression^15^. These regions are thought to function as genetic storage compartments, keeping certain genes silent. However, some LADs could be repositioned, moving either away from or toward the nuclear lamina in a cell-type-specific manner. This repositioning is critical for regulating gene expression, and any disruptions in this process can lead to altered transcriptional programs that contribute to *LMNA* cardiomyopathy^16^. Despite growing recognition of the nuclear lamina’s importance, much remains unclear regarding how these interactions influence cellular function and contribute to cardiac-specific disease caused by *LMNA* mutations.

In this study, we sought to explore the hypothesis that variants in the *LMNA* gene lead to the remodeling of chromatin and altered transcriptional programs, which may ultimately contribute to the pathophysiology of *LMNA* cardiomyopathy. To test this, we utilized both human induced pluripotent stem cells (hiPSC) reprogrammed from a patient with the *LMNA* single point mutation H222P mutation (*LMNA* p.H222P). For optimal comparison, we used an isogenic hiPSC control line corrected for the *LMNA* p.H222P mutation. We differentiated hiPSC-derived ventricular-like cardiomyocytes (hiPSC-CMs) in 2D cardiac sheets and employed a murine model expressing the same *LMNA* p.H222P mutation at the homozygous state (*Lmna*^H222P/H222P^)^17,18^. Through *in vitro* and *in vivo* functional, morphological, and transcriptomic high-throughput experiments, our findings revealed, for the first time, that *LMNA* variants cause distinct alterations in the chromosome spatial arrangement in hiPSC-CMs. These alterations were correlated with changes in the expression of genes associated with extracellular matrix organization. We reported specifically that the upregulation of Loxl2 plays a role in the development of *LMNA* cardiomyopathy. This work opens new insight into the pathogenesis of the diseases and could lead to a solution to treat cardiac remodeling.

## RESULTS

### Modeling LMNA cardiomyopathy in vitro

We employed the patient-specific *LMNA* p.H222P hiPSC line (*LMNA* H222P) and its isogenic control (*LMNA* corr.H222P) as previously published^19^. The patient-specific hiPSC were differentiated into hiPSC-CMs, enabling us to explore the cellular organization and contractile properties that may be altered in *LMNA* cardiomyopathy. Morphologically, the *LMNA* H222P hiPSC-CMs displayed a normal degree of sarcomere compaction (*LMNA* H222P, 1.495 ± 0.021 Lm; *LMNA* corr.H222P, 1.460 ± 0.026 Lm) **(Figure 1A, 1B)**. However, the cellular area of the mutated hiPSC-CMs was significantly smaller compared to *LMNA* corr.H222P hiPSC-CMs (*LMNA* H222P, 1212.5255 ± 101.805 Lm²; *LMNA* corr.H222P, 1353.934 ± 145.512 Lm²) **(Figure 1A, 1C)**. We observed reduced nuclear area (*LMNA* H222P, 87.400 ± 4.235 Lm²; *LMNA* corr.H222P, 99.215 ± 4.740 Lm²) and nuclear perimeter (*LMNA* H222P, 42.885 ± 0.922 Lm; *LMNA* corr.H222P, 46.303 ± 1.155 Lm) in mutated hiPSC-CMs compared to control cells **(Figure 1D, 1E)**. No differences in nuclear circularity or solidity (invaginations) were reported in *LMNA* H222P compared to *LMNA* corr.H222P hiPSC-CMs **(Figure 1F, 1G)**. We then evaluated the functional characteristics of the hiPSC-CMs, focusing on some key features of the excitation-contraction coupling. We monitored the intracellular calcium handling under electrical pacing in dissociated hiPSC-CMs using ratiometric dyes. The intracellular diastolic calcium level was significantly higher in the *LMNA* H222P hiPSC-CMs (*LMNA* H222P, 1.544 ± 0.026; *LMNA* corr.H222P, 1.388 ± 0.026) **(Figure 1H)**.

**Figure 1:**
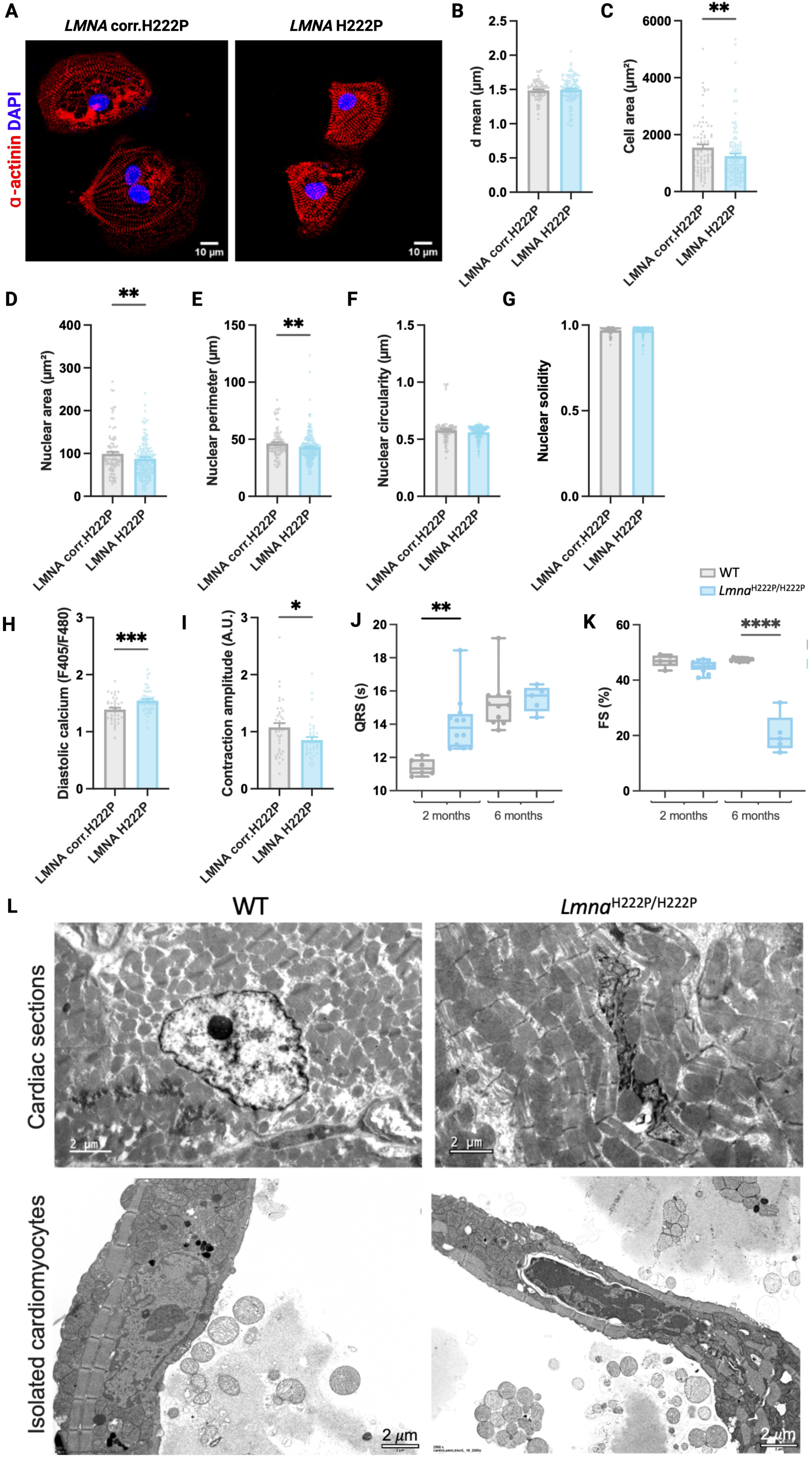
Mutated A-type lamins caused functional and structural impairments in both hiPSC-CMs and mice. **(A)** Representative immunostaining images of hiPSC-CMs. Sarcomeres were stained with an anti-α-actinin antibody (red) and the nuclear DNA was stained with DAPI (blue). Scale bar = 10 µm. **(B)** The mean distance between sarcomeres (dmean) in hiPSC-CMs (ns; n_H222P_= 3, n_corr.H222P_ = 3). **(C)** hiPSC-CMs cell area (** p = 0,0046; n_H222P_= 3, n_corr.H222P_= 3). **(D-G)** Nuclear structural parameters of hiPSC-CMs (N_H222P_= 2, n_H222P_=191, N_corr.H222P_= 2, n_corr.H222P_= 103). **(D)** Measure of nuclear area (** p = 0.0100). **(E)** Measure of the nuclear perimeter (** p = 0.0085). **(F)** Measure of nuclear circularity (ns). **(G)** Measure of nuclear solidity (ns). **(H)** Measurement of the ratio between the two emission wavelengths 405nm and 480nm under electrical pacing, reflecting the intracellular diastolic calcium concentration in *LMNA* H222P hiPSC-CMs (n_H222P_= 3, n_corr.H222P_= 2; *** p = 0,0008). **(I)** Measure of contraction amplitude of hiPSC-CMs under electrical pacing (* p= 0.0228; n_H222P_= 3, n_corr.H222P_= 3). **(J)** QRS time duration measurement in *Lmna^H^*^222^*^P/H^*^222^*^P^ mice (*\*\* p= 0.0415; one-way ANOVA, mean ± SEM; n_WT-2m_= 6, n_Lmna-2m_ = 12, n_WT-6m_ = 11, n_Lmna-6m_ = 5). **(K)** Fractional shortening measurement by echocardiography in *Lmna^H^*^222^*^P/H^*^222^*^P^* mice (**** p < 0.0001, ANOVA statistic test, mean ± SEM; n_WT-2m_= 6, n_Lmna-2m_ = 12, n_WT-6m_ = 11, n_Lmna-6m_ = 5). **(L)** Representative images from electron microscopy of cell nuclei of myocytes from heart slices (top panel) and isolated *Lmna^H^*^222^*^P/H^*^222^*^P^* mice myocytes, aged 6 months (bottom panel). Scale bar = 2 µm. All functional experiments on hiPSC-CMs were analyzed using a Mann-Whitney statistical test, mean ± SEM.

Elevated diastolic calcium levels are often associated with cardiac arrhythmias and contractile dysfunction, both of which are common features of DCM^20^. The abnormal calcium handling could result from defects in calcium signaling pathways, which are essential for cardiac muscle contraction and relaxation^21,22^. Using video-edge capture, we measured the contractile properties under electrical pacing in dissociated hiPSC-CMs. We consistently observed disrupted 2D cardiac sheets with the *LMNA* H222P hiPSC-CMs, suggesting high tissue fragility. We found that *LMNA* H222P hiPSC-CMs exhibited reduced contraction amplitude (hypocontractility) (*LMNA* H222P, 0.853 ± 0.053 A.U.; *LMNA* corr.H222P, 1.075 ± 0.077 A.U.) compared to *LMNA* corr.H222P hiPSC-CMs **(Figure 1I)**. We monitored the proportion of arrhythmic events in hiPSC-CMs by assessing the heterogeneity of their contractions. No significant difference was observed between the *LMNA* H222P and *LMNA* corr.H222P, but the heterogeneity of contraction tended to be higher in the mutant than in the control (*LMNA* H222P, 1.111 ± 0.103; *LMNA* corr.H222P, 1.278 ± 0.116). hiPSC-CMs **(Figure S1A)**. The contraction (*LMNA* H222P, 73.534 ± 4.213; *LMNA* corr.H222P, 138.830 ± 7.109) and diastolic (*LMNA* H222P, 191.291 ± 13.692; *LMNA* corr.H222P 299.471 ± 12.685) times were increased in *LMNA* H222P compared to *LMNA* corr.H222P **(Figure S1B, S1C)**.

To confirm our *in vitro* findings in an integrated model, we examined the *Lmna^H^*^222^*^P/H^*^222^*^P^* mouse model at 2 and 5 months old, corresponding to pre- and post-cardiac symptomatic stages^18^. *Lmna^H^*^222^*^P/H^*^222^*^P^* mice exhibited an abnormal electrocardiogram profile, which was characterized by a prolonged QRS interval without reaching significativity (*Lmna^H^*^222^*^P/H^*^222^*^P^*, 2 months 14.028 ± 0.482 s, 5 months 15.534 ± 0.345 s; WT, 2 months 11.419 ± 0.203 s, 5 months 15.302 ± 0.450 s) **(Figure 1J)**, which reflects cardiac conduction defects. The contractile function was also compromised, as indicated by a decreased fractional shortening (*Lmna^H^*^222^*^P/H^*^222^*^P^*, 2 months 44.848 ± 0.571 %, 5 months 20.536 ± 3.065 %; WT, 2 months 46.960 ± 0.999 %, 5 months 47.405 ± 0.180 %) **(Figure 1K)**. As a hallmark of *LMNA* cardiomyopathy, we investigated the nuclear morphology in heart tissue sections and isolated cardiomyocytes from *Lmna^H^*^222^*^P/H^*^222^*^P^*mice. The cardiomyocytes displayed elongated nuclei and nuclear envelope abnormalities, including nuclear envelope detachment, predominantly observed in isolated cardiomyocytes **(Figure 1L).** Collectively, these findings highlight that complementary model systems, including patient-derived hiPSCs and mice carrying the same disease-relevant *LMNA* mutation, provide a robust platform for cross-species, in-depth exploration of *LMNA* cardiomyopathy pathogenesis mechanisms *in vitro* and *in vivo*.

### LMNA H222P hiPSC-CMs exhibit altered chromosome positioning and extracellular matrix-related gene expression

In eukaryotic cells, chromosomes occupy distinct territories within the nucleus, which is important for genome compaction and regulation^23^. To assess the impact of the *LMNA* p.H222P mutation on chromosome organization, we performed chromosome painting and quantified their distribution in hiPSC-CMs **(Figure 2A, 2B)**. We reported that the largest chromosomes (e.g., chr1, chr2, chr3) were significantly located towards the nuclear periphery in the *LMNA* H222P compared to *LMNA* corr.H222P hiPSC-CMs **(Figure 2C)**. A-type lamins are essential for genome organization by regulating the distribution of LADs, which in turn influences gene expression^24–27^. We next sought to identify abnormal expression of genes involved in the development of cardiomyopathy caused by *LMNA* mutation using a bulk RNA-seq in hiPSC-CMs. This analysis yielded 607 differentially expressed genes (DEGs) between *LMNA* H222P and *LMNA* corr.H222P hiPSC-CMs **(Figure 3A, Figure S2A, S2B)**. We reported that the chromosomes with a spatial remodeling **(Figure 2C)** exhibited a higher proportion of DEGs **(Figure 3B)**. We next performed gene ontology analysis to identify the biological processes associated with the DEGs in *LMNA* H222P hiPSC-CMs. These DEGs were primarily involved in ECM organization and developmental processes **(Figure 3C)**. We observed that DEGs related to ECM organization were mostly present in chromosome 1 **(Figure 3D)**. To confirm the relevance of these findings in fully developed diseased hearts, we conducted a genome-wide RNA expression analysis on heart tissues from *Lmna^H^*^222^*^P/H^*^222^*^P^* mice. This analysis revealed 3,487 DEGs **(Figure 3E, Figure S2C, S2D)** primarily associated with mitochondria metabolism and ECM organization **(Figure 3F)**. The dysregulation of these genes could contribute to pathological processes such as fibrotic remodeling and electrical conduction defects, both of which are hallmarks of DCM^28^. A comparison of biological processes linked to the DEGs in both *in vitro* and *in vivo* models revealed a common focus on ECM structure, organization, and remodeling **(Figure 3G)**.

**Figure 2:**
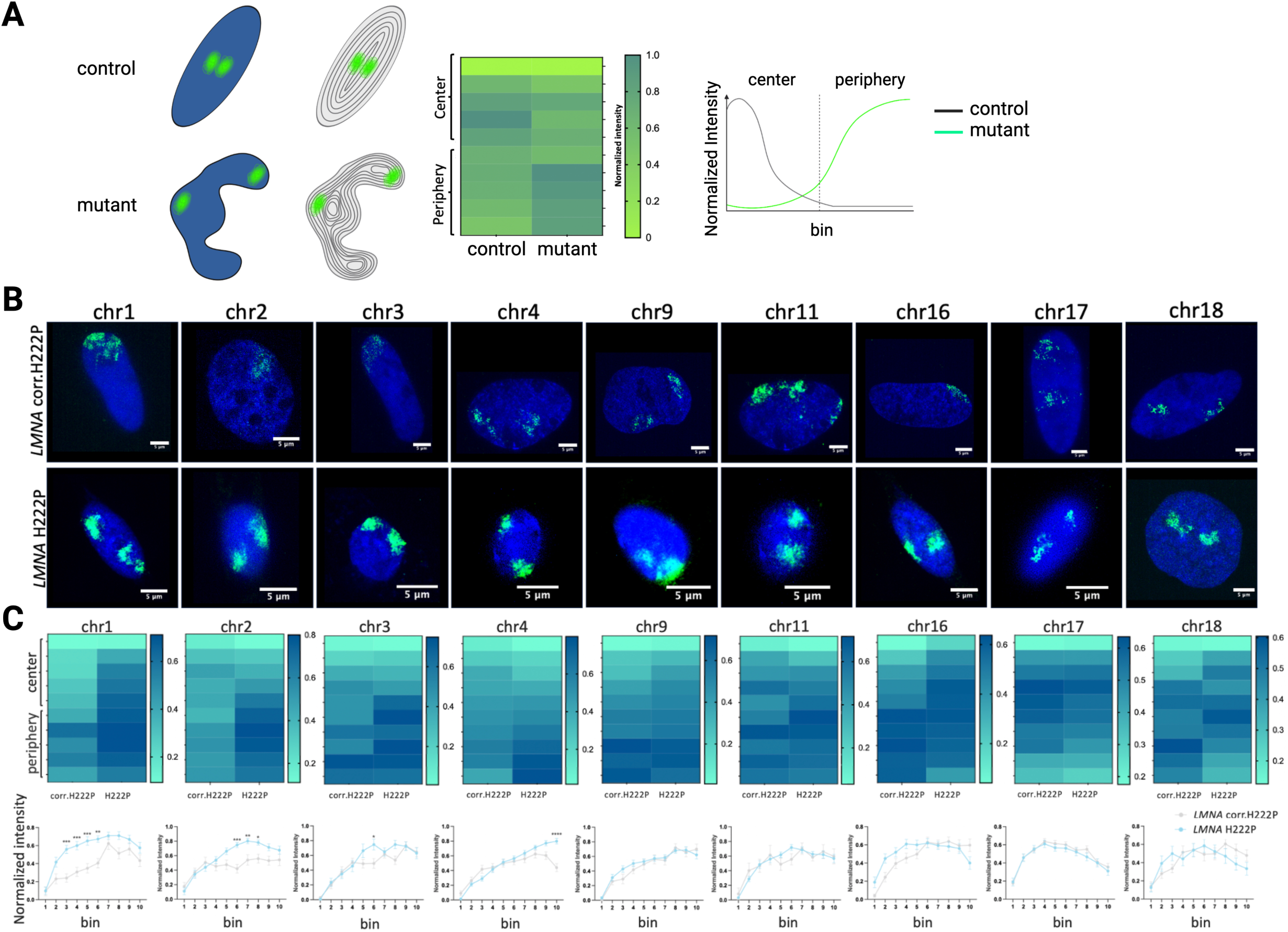
Mutated A-type lamins altered the chromosome spatial organization in hiPSC-CMs. **(A)** Schematic pipeline of chromosome painting analysis. DAPI channel images were binarized to create nuclear masks which were divided into ten concentric rings (bins). Bins 1-5 correspond to the nuclear center and bins 6 to 10 correspond to the nuclear periphery. 488 and 561 channel images, corresponding to chromosome probes, were superposed to the corresponding nuclei mask to quantify chromosome positions. The results are represented with heatmaps or graphs. **(B)** Representative images of chromosome painting experiments. Scale bar = 5 µm. **(C)** Heatmaps and graphs representing the results from the chromosome painting experiment (chr1, *** p= 0.0001, *** p= 0.0006, *** p= 0.0003, ** p= 0.0025; chr2, *** p= 0.0004, ** p= 0.01, * p= 0.0495; chr3, * p= 0.0385; two-way ANOVA statistic test, mean ± SEM; n_H222P_= 2, n_corr.H222P_= 2).

**Figure 3:**
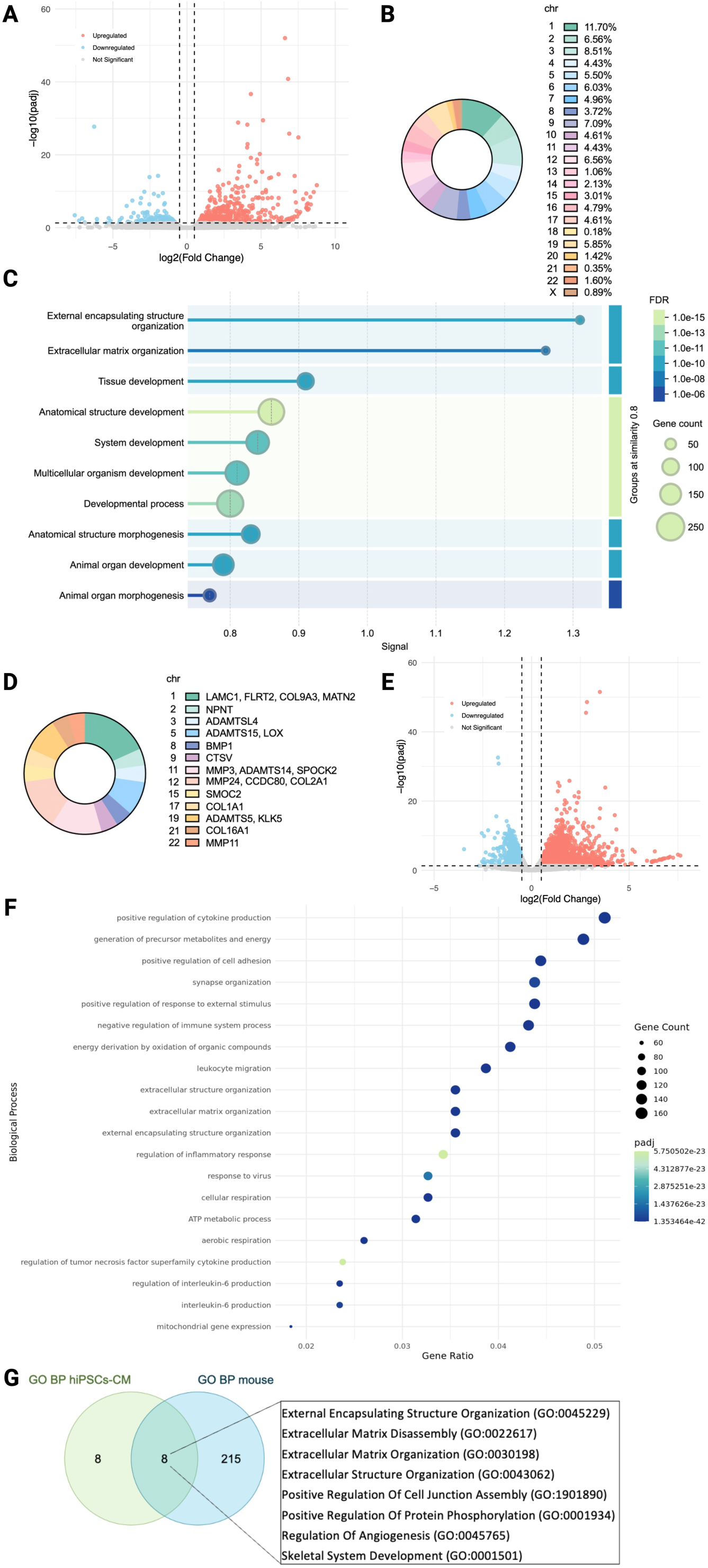
Mutated A-type lamins lead to gene expression dysregulation in both hiPSC-CMs and mice. **(A)** Volcano plot representing DEGs between *LMNA* H222P and *LMNA* corr.H222P hiPSC-CMs. **(B)** Graphical representation of DEGs percentage per chromosome in hiPSC-CMs. **(C)** Graphical representation of biological processes enrichment associated with DEGs in hiPSC-CMs. Generated with STRING v12.0. **(D)** Graphical representation of ECM-related DEGs per chromosome in hiPSC-CMs. **(E)** Volcano plot representing DEGs between *Lmna^H^*^222^*^P/H^*^222^*^P^* and WT mice. **(F)** Graphical representation of first 20 biological processes associated with DEGs in mice. **(G)** Venn diagram and table recapitulating the common biological processes associated with DEGs in common between hiPSC-CMs and mice. (n_H222P_= 2, n_corr.H222P_= 3, n_Lmna_= 3, n_WT_= 2; threshold, padj 0.05, log2FC 0.5).

### Loxl2 mediates LMNA cardiomyopathy in vivo

The role of ECM remodeling in *LMNA* cardiomyopathy remains underexplored. However, we observed significant fibrosis development in hearts from *Lmna^H^*^222^*^P/H^*^222^*^P^* mice compared to WT mice as demonstrated by histology and electron microscopy **(Figure 4A, 4B)**. These findings were further supported by elevated *Col1a2* and *Col3a1* mRNA expression in *Lmna^H^*^222^*^P/H^*^222^*^P^* mice compared to WT mice **(Figure 4C, 4D)**, both of which are established biomarkers of cardiac diseases, including heart failure^29^.

**Figure 4:**
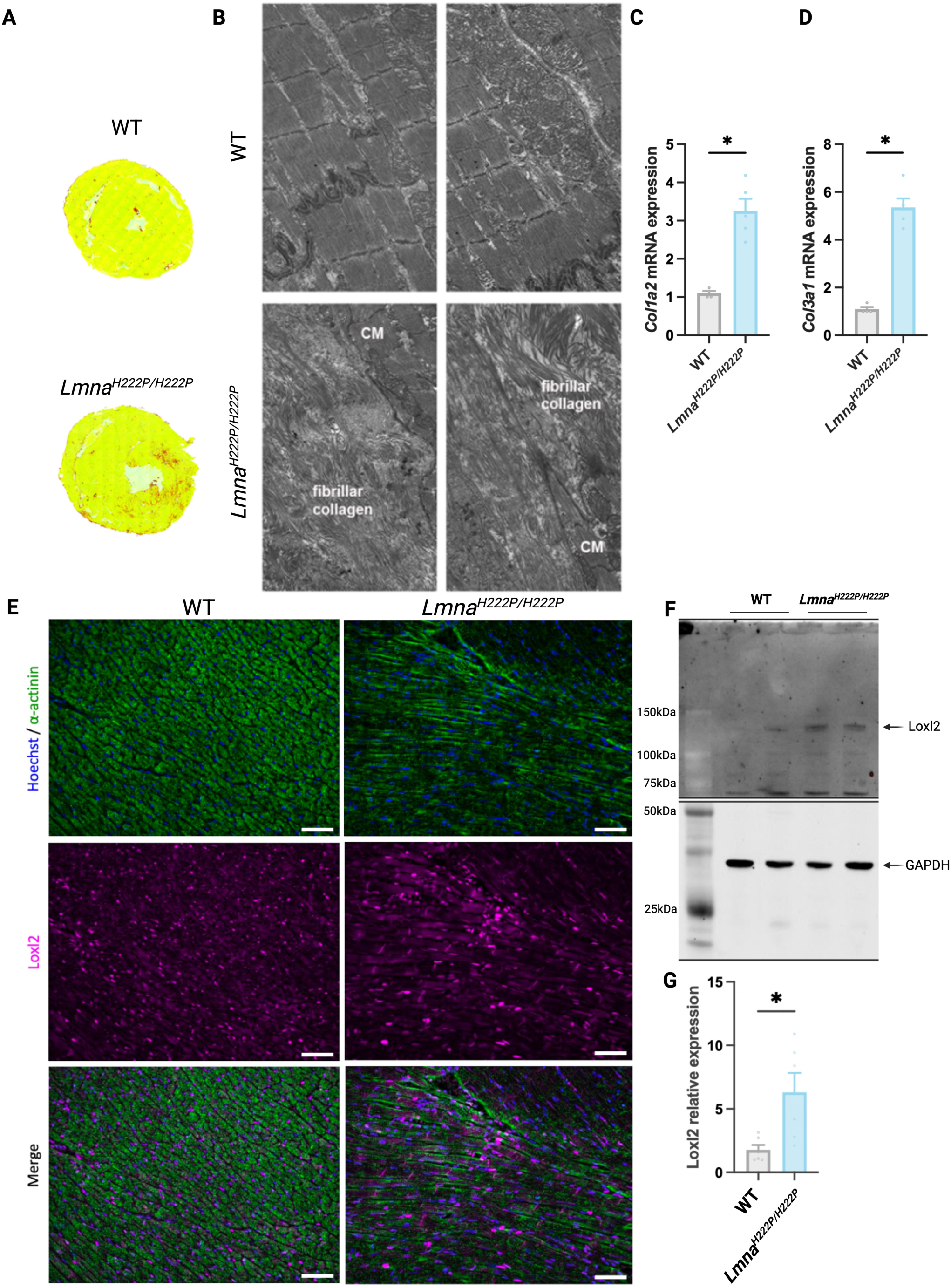
Mutated A-type lamins triggered Loxl2 protein expression increase in mice. **(A)** Transverse cardiac sections of *Lmna^H^*^222^*^P/H^*^222^*^P^* and WT mice colored with Sirius red. **(B)** Representative electron microscopy images of fibrous collagen in WT (top) and *Lmna^H^*^222^*^P/H^*^222^*^P^* (bottom) mice. **(C,D)** mRNA expression of **(C)** *Col1a2* and **(D)** *Col3a1* in *Lmna^H^*^222^*^P/H^*^222^*^P^*and WT mice (* p= 0.0159; Mann-Whitney statistic test, mean ± SEM). **(E)** Representative immunostaining images of Loxl2 protein expression in mice (magenta). Sarcomeric actin was stained in green and nuclear DNA in blue. Scale bar= 100µm **(F)** Immunoblot of Loxl2 protein expression. GAPDH protein expression was used to quantify protein expression (n_Lmna_= 2, n_WT_= 2). **(G)** Quantification of Loxl2 protein expression (* p= 0.0161; Unpaired t-test, mean ± SEM).

We identified several key dysregulated genes related to ECM establishment, remodeling, and fibrosis in hearts from *Lmna^H^*^222^*^P/H^*^222^*^P^* compared to wild-type (WT) mice. Among these genes, we focused on *Loxl2* **(Figure 4A)**, as it has been previously reported to be upregulated in failing hearts from mice and linked to increased interstitial fibrosis and cardiac dysfunction^30^. Bulk RNA-seq data confirmed the upregulation of *Loxl2* in the hearts from *Lmna^H^*^222^*^P/H^*^222^*^P^* mice compared to WT mice (padj 4.035E-05 log2FC 1.869) **(Figure 4A)**, with a similar trend observed in *LMNA* H222P hiPSC-CMs (pval 0.0122, padj 0.260, log2FC 1.296). We next sought to correlate changes in *Loxl2* gene expression with Loxl2 protein overexpression *in vivo*. We performed immunostaining on cardiac sections from *Lmna^H^*^222^*^P/H^*^222^*^P^* mice. Loxl2 was upregulated in the mutated mice compared to WT mice **(Figure 4B)**. These findings were further validated by immunoblot experiments **(Figure 4C, 4D)**.

To further decipher the role of Loxl2 in the development of *LMNA* cardiomyopathy, we performed antibody-mediated Loxl2 inhibition in *Lmna^H^*^222^*^P/H^*^222^*^P^* mice. Cardiac function and morphology were evaluated by echocardiography before treatment and one month post-treatment to assess the impact of Loxl2 depletion **(Figure 5A)**. This approach allowed us to quantify changes directly attributable to Loxl2 inhibition and its potential involvement in *LMNA* cardiomyopathy pathogenesis. We measured the left ventricular end diameter in systole (LVEDs) **(Figure 5B, 5C)** and diastole (LVEDd) **(Figure 5 D, 5E)**, the ejection fraction (EF) **(Figure 5F, 5G)** and the fractional shortening (FS) **(Figure 5H, 5I)**. After one month of treatment with Loxl2 antibody, we observed preservation of cardiac function and prevention of left ventricular dilatation in treated *Lmna^H^*^222^*^P/H^*^222^*^P^* mice (Lmna+Loxl2) compared to untreated *Lmna^H^*^222^*^P/H^*^222^*^P^* mice (Lmna no treat.) **(Figure 5B-5I)**. While untreated *Lmna^H^*^222^*^P/H^*^222^*^P^* mice exhibited progressive cardiac dysfunction and left ventricular dilatation between 4 and 5 months, these parameters remained stable in treated *Lmna^H^*^222^*^P/H^*^222^*^P^* mice during the same period **(Figure 5B-5I)**. Notably, each treated *Lmna^H^*^222^*^P/H^*^222^*^P^* mice showed a stabilization of these parameters over time between 4 and 5 months, in contrast to their deterioration observed during the same period in untreated *Lmna^H^*^222^*^P/H^*^222^*^P^* mice **(Figure 5C, 5E, 5G, 5J)**. Together, these results suggest that inhibition of Loxl2 functionally preserves cardiac function in a mouse model of *LMNA* cardiomyopathy.

**Figure 5:**
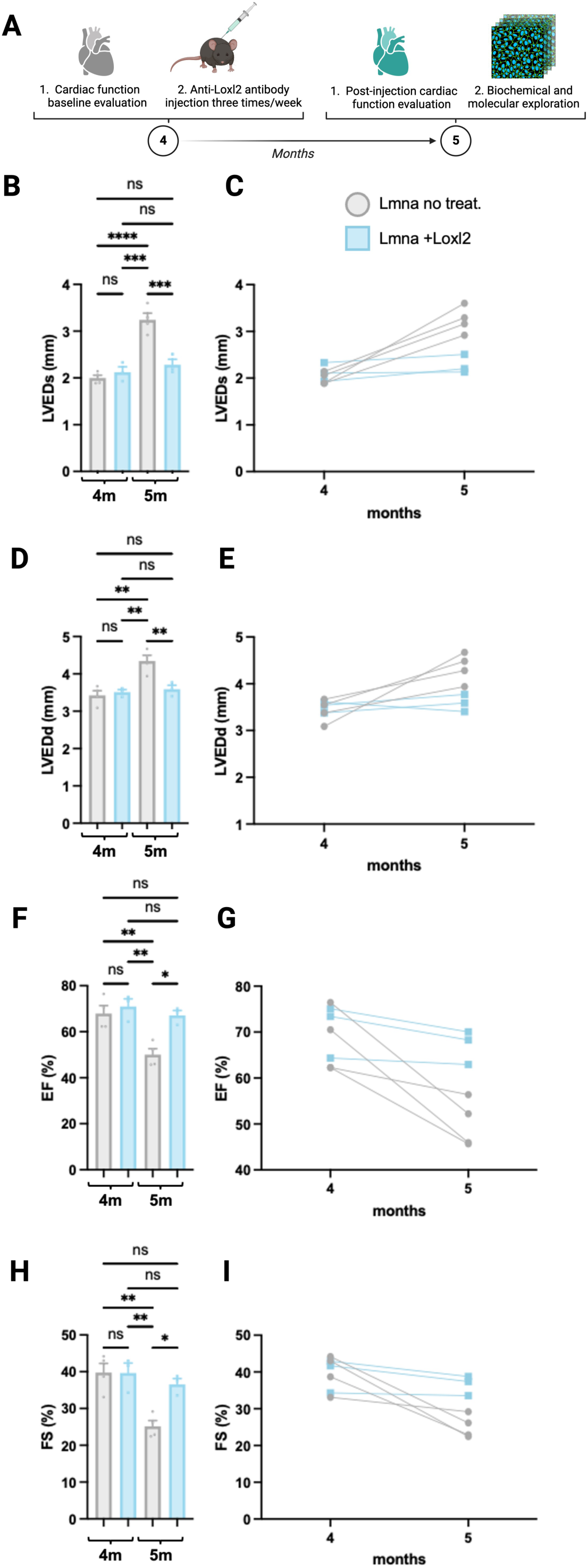
Loxl2 inhibition treatment preserved cardiac function in *Lmna^H^*^222^*^P/H^*^222^*^P^* mice. **(A)** Schematic representation of Loxl2 mice treatment. Echocardiography and electrocardiography were performed on *Lmna^H^*^222^*^P/H^*^222^*^P^* mice before Loxl2 treatment. During one month, every two days, half of the mice received an IP injection with an anti-Loxl2 antibody treatment. After one month, electrocardiography and echocardiography were performed to evaluate the effect of Loxl2 on cardiac function and morphology. Hearts were taken for further biochemical and molecular analysis. Generated with BioRender. **(B-I)** Echocardiography of *Lmna^H^*^222^*^P/H^*^222^*^P^*mice with or without treatment, before and after treatment. **(B,C)** Measure of the left ventricular end diameter in systole (LVEDs) (**** p <0.0001, *** p= 0.0002, *** p= 0.0007) and **(D,E)** in diastole (LVEDd) (** p= 0.0014, ** p= 0.0050, ** p= 0.0095), reflecting the cardiac structure. **(F,G)** Measure of the ejection fraction (EF) (** p= 0.0055, ** p= 0.0031, * p= 0.0121) and **(H,I)** fractional shortening (FS) (** p= 0.0028, ** p= 0.0051, * p= 0.0244) reflecting the cardiac function. (n_Lmna+Loxl2_= 3, n_Lmna no treat._= 4). All the echocardiography parameters were analyzed using a one-way ANOVA statistic test, mean ± SEM. no treat., no treatment with the anti-Loxl2 antibody; +Loxl2, treated with the anti-Loxl2 antibody.

## DISCUSSION

In this study, we successfully modeled *LMNA* cardiomyopathy *in vitro* using patient-specific *LMNA* p.H222P hiPSC lines (*LMNA* H222P) and its isogenic control (*LMNA* corr.H222P), complemented by an *in vivo Lmna*^H222P/H222P^ mouse model. Our findings provide critical insights into the cellular, molecular, and functional alterations associated with *LMNA* cardiomyopathy, highlighting the role of nuclear architecture, intracellular calcium handling, contractile dysfunction, and ECM remodeling in disease pathogenesis.

The differentiation of patient-specific hiPSC-CMs allowed us to investigate the morphological and functional consequences of the *LMNA* p.H222P mutation. While the *LMNA* H222P hiPSC-CMs exhibited normal sarcomere compaction, they were significantly smaller than *LMNA* corr.H222P hiPSC-CMs. Additionally, these cells displayed abnormal nuclear morphology, including altered nuclear area and perimeter, consistent with previous studies demonstrating that *LMNA* mutations disrupt nuclear envelope integrity^24,27^. These nuclear abnormalities were further corroborated in the *Lmna^H^*^222^*^P/H^*^222^*^P^* mouse model, where cardiomyocytes exhibited elongated nuclei and nuclear envelope detachment. These findings underscore the role of A-type lamins in maintaining nuclear structure and suggest that nuclear defects may contribute to impaired cellular function by disrupting chromatin organization and gene expression.

Functional analyses of *LMNA* H222P hiPSC-CMs revealed elevated intracellular diastolic calcium levels and hypocontractility, both of which are hallmarks of DCM. Interestingly, similar defects have been observed in patient-derived Duchenne muscular dystrophy hiPSC-CMs^31^, suggesting common defects in *LMNA* and *DMD* cardiomyocytes affecting the excitation-contraction coupling. Abnormal calcium handling, likely resulting from defects in calcium signaling pathways critical for cardiac muscle contraction and relaxation^21,22^, may underlie the arrhythmogenic and contractile dysfunction observed in *LMNA* cardiomyopathy. We found no evidence of arrhythmic events at rest in dissociated *LMNA* H222P hiPSC-CMs when measuring the contractile properties by video-edge capture. It should be noted that monitoring single-cell contractions might not accurately reflect the defects occurring in cardiac tissue. Notably, we could not measure these properties on the integrated 2D cardiac sheet due to the consistent early-stage disruption of the monolayer observed in LMNA H222P hiPSC-CMs, suggesting high tissue fragility likely leading to conductivity defects. We observed higher contraction and diastolic times in *LMNA* H222P hiPSC-CMs, which could indicate damaged and weaker cardiomyocytes that appeared smaller. The contraction heterogeneity was similar between *LMNA* H222P and *LMNA* corr.H222P hiPSC-CMs, although a tendency towards greater heterogeneity might suggest an increased likelihood of arrhythmic events in the mutated cells. We also monitored the electrophysiological features of the *Lmna^H^*^222^*^P/H^*^222^*^P^* mouse model *in vivo*. The *in vitro* findings were validated in the *Lmna^H^*^222^*^P/H^*^222^*^P^*mouse model, which exhibited prolonged QRS intervals and decreased fractional shortening, indicative of compromised cardiac conduction and contractile function. Prolonged QRS intervals are a significant clinical feature observed in patients with *LMNA* cardiomyopathy and are indicative of impaired cardiac conduction. Together, these results highlight the critical role of intracellular calcium mishandling and hypocontractility in the pathogenesis of *LMNA* cardiomyopathy.

Our results indicated that the *LMNA* p.H222P mutation also significantly alters chromosome positioning in hiPSC-CMs, with larger chromosomes (e.g., chr1, chr2, chr3) localized more toward the nuclear periphery. This spatial remodeling may reflect disrupted interactions between lamin A/C and LADs, which are essential for genome organization and gene regulation^23,24^. RNA-seq analysis identified dysregulated genes mainly involved in ECM organization and development in *LMNA* H222P compared to *LMNA* corr.H222P hiPSC-CMs. Similar dysregulation of ECM-related genes was observed in heart tissue from *Lmna^H^*^222^*^P/H^*^222^*^P^* mice, suggesting a conserved pathogenic mechanism across species. The overlap in DEGs related to ECM structure and remodeling in both models highlights the central role of ECM dysregulation in *LMNA* cardiomyopathy.

We further established that *LMNA* cardiomyopathy is associated with impaired ECM gene regulation, as evidenced by both patient-derived *in vitro* and murine *in vivo* models. The proper functioning of the heart relies on the interplay of numerous factors, whose dysregulation can contribute to cardiac pathogenesis, including *LMNA* cardiomyopathy. Among these, the ECM plays a central role in maintaining cardiac homeostasis and is a key player in both physiological and pathological states. The cardiac ECM is composed of a diverse array of proteins, including collagens (collagen I, III, and IV), non-collagenous glycoproteins (fibronectin, laminin), proteoglycans, glucosaminoglycans, and elastin^32^. In the context of cardiac disease, such as heart failure, or during injury, such as myocardial infarction, physiopathological stress triggers ECM remodeling. This process often involves excessive ECM deposition by cardiac fibroblasts (fibrosis) or alterations in their biochemical composition, leading to increased myocardial stiffness and aberrant signal transmission to cardiomyocytes. These changes ultimately result in impaired contractility^33^. Fibrosis is particularly critical in DCM, where it significantly contributes to cardiac dysfunction and exacerbates the severity of the clinical phenotype. Together, these findings highlight the importance of ECM remodeling in the pathogenesis of *LMNA* cardiomyopathy and suggest that targeting ECM dysregulation may offer therapeutic potential for this devastating disease. Myocardial septal fibrosis, a hallmark of *LMNA* cardiomyopathy^5–7^, is linked to conduction abnormalities like atrioventricular block and PR-interval prolongation^34^. The TGF-β/Smad pathway drives fibrosis by promoting fibroblast proliferation, myofibroblast differentiation, and ECM protein deposition^35^ while suppressing matrix-degrading enzymes^36^. Upregulation of YY1 to enhance BMP7 and limit CTGF (CCN2) reduces fibrosis and improves cardiac function in *LMNA* cardiomyopathy^37^. TGF-β/Smad also activates ERK1/2 signaling, which exacerbates fibrosis and ventricular dysfunction in *Lmna^H^*^222^*^P/H^*^222^*^P^* mice^38–40^. These findings highlight the critical role of ECM regulation in *LMNA* cardiomyopathy pathogenesis. One key gene identified here was Loxl2, which was upregulated in the *Lmna^H^*^222^*^P/H^*^222^*^P^* compared to WT mice. These results are consistent with the role of Loxl2 in promoting fibrosis and cardiac dysfunction^30^. Immunostaining and immunoblot experiments confirmed Loxl2 overexpression in *Lmna^H^*^222^*^P/H^*^222^*^P^*mice, and antibody-mediated inhibition of Loxl2 preserved cardiac function and morphology in treated mice. These findings suggest that Loxl2 contributes to the fibrotic remodeling and contractile dysfunction characteristic of *LMNA* cardiomyopathy and may represent a potential therapeutic target. It has been shown that Loxl2 is involved in crosslinking collagen and elastin^41^, which is crucial for ECM stability. Importantly, Loxl2 has been identified as a biomarker of heart failure^30^. In mouse models of heart failure^30^ or diabetic cardiomyopathy^42^, upregulation of Loxl2 promotes interstitial fibrosis, leading to cardiac dysfunction. Notably, inhibition of Loxl2, either through antibody-mediated blockade, genetic deletion, or siRNA, significantly mitigates interstitial fibrosis and improves cardiac function^30,42^. Loxl2 is notably upregulated by TGF-β, inflammation, mechanical stimuli, and hypoxia in endothelial and vascular smooth muscle cells^30,43–45^. These factors are critical to cardiac homeostasis, and pathogenesis, suggesting they could similarly contribute to Loxl2 upregulation in the heart. Understanding these mechanisms provides insight into Loxl2’s role in cardiac remodeling and its potential as a therapeutic target.

Our findings highlight the importance of ECM regulation in cardiac homeostasis and underscore the therapeutic potential of targeting ECM components, such as Loxl2, in mitigating fibrosis and preserving cardiac function in DCM. While Loxl2 has been involved in other conditions, including heart failure^30^ and diabetic cardiomyopathy^42^, this study marks the first time Loxl2 has been associated with *LMNA* cardiomyopathy. Consequently, Loxl2 and its gene regulatory network emerge as promising therapeutic targets for enhancing cardiac function in *LMNA* cardiomyopathy.

While our study provides valuable insights, certain limitations should be acknowledged. The patient-derived cardiomyocyte model, while powerful, may not fully recapitulate the complexity of adult cardiomyocytes or the chronic disease state observed in patients. We used a corrected isogenic control hiPSC line to evaluate the consequences of the *LMNA* p.H222P mutation. This strategy is currently the best for assessing the genotype/phenotype relationship with minimal polymorphism. However, isogenic control hiPSC lines may not always behave like unaffected healthy control hiPSC lines. Additional healthy control lines would strengthen our results. Additionally, the *in vivo* effects of Loxl2 inhibition were evaluated over a relatively short period, and long-term studies are needed to assess sustained therapeutic benefits. Further investigation into the interplay between nuclear architecture, calcium handling, and ECM remodeling will be essential to fully elucidate the mechanisms driving *LMNA* cardiomyopathy.

In conclusion, our findings underscore the multifaceted nature of *LMNA* cardiomyopathy, involving disruptions in nuclear structure, calcium homeostasis, contractile function, and ECM remodeling. The identification of Loxl2 as a potential therapeutic target offers hope for developing effective treatments for this devastating disease. The use of patient-specific hiPSC models, combined with *in vivo* validation, underscores the potential of these models to unravel disease mechanisms and identify therapeutic targets. Future studies should focus on further elucidating the pathways involved in ECM remodeling and exploring targeted therapies to mitigate the effects of *LMNA* mutations on cardiac function.

## MATERIAL AND METHODS

### Ethics statement

The study adhered to the principles of the Declaration of Helsinki as previously reported^19^.

### hiPSC lines

We used the patient-specific *LMNA* p.H222P hiPSC line (*LMNA* H222P) and its isogenic control (*LMNA* corr.H222P) as previously published^19^. All hiPSC lines were maintained as previously reported^46^.

### Cardiac differentiation

Once hiPSC reached 90% confluency, 2D sandwich-based differentiation into ventricular-like hiPSC-CMs followed the Wnt pathway activation/inhibition protocol as described in^47^. On day 9, the medium was changed to RPMI 1640-B27. From day 9, hiPSC-CMs were further maturated with 100nM 3,3’,5 triodo-L-thyronine (T3) (Sigma-Aldrich, Cat#T2877) and 0.5mM N6,2′-O-Dibutyryladenosine 3′,5′-cyclic monophosphate sodium salt (dAMPc) (Sigma-Aldrich, Cat#D0627)^48,49^. By day 20, hiPSC-CMs were purified using 4mM sodium L-lactate (Sigma-Aldrich, Cat#L7022) in Silac medium (Gibco, Cat#A2494201), according to ^50^. All the experiments were performed between day 30 to day 35 of cardiac differentiation.

### Diastolic calcium level quantification

Diastolic calcium levels were assessed using the ratiometric fluorescent Ca^2+^ sensor dye Indo-1-AM (Molecular Probes, Cat#I1223). Enzymatically-dissociated hiPSC-CMs were incubated for 25 min at 37°C with 2 mM Indo-1-AM, in the Tyrode solution. The diastolic calcium level was measured using Ionoptix acquisition system FSI700 fluorescence with PMT400, under electrical pacing (1Hz, 20V, 5ms duration). All ratios were measured using Ionoptix Software.

### Measurement of contractile properties by video-edge capture

Enzymatically-dissociated hiPSC-CMs were seeded in microscopy dishes (Ibidi, Cat#81156) containing 2 mL of 1.8 mM to measure the contractile parameters under electrical pacing (1Hz, 20V, 5ms duration), using microscopy video-edge capture (63 frames/sec, 25 s/positions, 15 ms/image), with x63 oil objective of Zeiss LSM900 and Zen Software at 37°C. Acquisitions were analyzed using a MatLab homemade script as published before^46,51,52^.

### Cellular morphology assessment

Enzymatically-dissociated hiPSC-CMs were seeded in microscopy dishes (Ibidi, Cat#81156). Cells were fixed with 4% PFA solution for 15 min at RT, permeabilized using 2 mL of permeabilization solution (1X PBS + 0.01% Tritton 100X) for 10 min at RT, and incubated with 1 mL of blocking solution (1X PBS + 1% BSA) for 30 min at RT. For immunostaining, cells were incubated 1 h at RT or O/N at 4°C with 500 mL of a solution of primary antibodies (1X PBS + 0.1% BSA). Cells were incubated 2 h at RT protected from light, with secondary antibodies solution (1X PBS + 0.01% BSA). Images acquisition was performed using X63 oil objective of Zeiss LSM900 microscope and Zen Software. To measure the cell morphology and organizational parameters, images were analyzed using MorphoScript under MatLab as described in ^53^.

### Nuclear morphology assessment

Enzymatically-dissociated hiPSC-CMs were fixed, permeabilized, and then marked with DAPI, following the same protocol as in the section “Cellular Morphology Assessment”. The images were acquired using a Nikon Ti2 microscope, equipped with a motorized plate and an x100 oil objective with a Prime 95B Scientific CMsOS (sCMsOS) camera. The microscope was controlled by MetaMorph 7.10 (Molecular Devices) software with a Pixel resolution of 0.11μm/px at 16 bits. Images were pre-treated with FIJI/ImageJ software. The nucleus was segmented using CellPose^54^ v2.2.2 with a GPU NVIDIA GeForce RTX 3080 Ti, using the performed model cyto2 and a diameter fixed at 120 pixels. Segmentation masks were imported in QuPath^55^ v0.3.2, where nuclear parameters were measured.

### Murine Lmna^H222P/H222P^ model

Homozygous males *Lmna*^H222P/H222P^, previously characterized in ^18,56^ were used for this study. Mice were provided standard chow and water ad libitum and maintained in a disease-free barrier facility under controlled conditions: 12 h light/dark cycle, a temperature of 22 °C, and 55% relative humidity. All animal experiments were approved by the French Ministry of Higher Education and Research and conducted at the Center for Research in Myology in compliance with the European Directive 2010/63/EU on the protection of animals used for scientific purposes.

### Functional and morphological heart exploration

Mice were anesthetized with 0.75% isoflurane in O_2_ and placed on a heating pad (28 °C). Transthoracic echocardiography was performed using a Vivid 7 Dimension/Vivid7 PRO ultrasound with an 11MHz transducer applied to the chest wall. Cardiac ventricular dimensions and fractional shortening were measured in 2D mode and M-mode three times for the number of animals indicated. A ‘blinded’ echo-cardiographer, unaware of the genotype and the treatment, performed the examinations.

Electrocardiograms were recorded from mice using the non-invasive ecgTUNNEL (Emka Technologies) with minimal filtering. Waveforms were recorded using Iox Software and intervals were measured manually with ECG Auto software. The electrocardiographer was blinded to the mouse genotype.

### Chromosome painting assay

Chromosome territories were visualized using the chromosome fluorescent *in situ* hybridization (FISH) approach, known as chromosome painting. The probes used were whole-chromosome and coupled with green (505 nm) or orange (552 nm) fluorophores (MetaSystems Probes, Cat #XCyting Chromosome Paint). The chromosome painting experiment was performed according to a combination of manufacturer’s and published protocols^57^. Briefly, hiPSC-CMs were seeded in glass coverslips, and then fixed with 4% PFA 20 min at RT, permeabilized with 0.05% Triton 100X 20 min at RT, and depurinated with 0.1M HCl 5 min at RT. Cells were washed twice with 2X SSC, 5 min at RT, and with 50% formamide/4X SSC at 37°C, 4 h in a humidified chamber. Cells and probes were mixed on a slide, sealed with rubber cement (Talens, Cat #95306500), and denatured at 75°C for 2 min. The hybridization step was performed at 37°C, O/N in a humidified chamber. Cells were washed with 0.4X SSC at 75°C, 2 min, 2X SSC + 0.05% Tween-20, 30 sec, and H2O to avoid crystallization. Cells were counterstained with DAPI for 10min at RT, light-protected, and a coverslip was mounted in a glass slide. The chromosome organization was visualized using a Nikon Ti2 microscope coupled with a live super-resolution module (Live-SR 3D, Gataca Systems), equipped with a motorized plate, and coupled with x100 oil objective with Prime 95B Scientific CMsOS camera (sCMsOS). The acquisition was realized using MetaMorph 7.10 (Molecular Devices) software with a Pixel resolution of 0.0658 μm/px at 16-bits. Nuclei were manually cut out before acquisition to avoid bleaching on other nuclei. A pipeline was developed to measure the chromosome distribution into the nucleus Nucleus masks were obtained using QuPath^55^ and binarized with ImageJ/Fiji. An algorithm was implemented to QuPath^55^ to divide each detected binarized nucleus mask into 10 bins. The bins from 1 to 5 correspond to the nuclear periphery whereas bins from 6 to 10 correspond to the nuclear center. This approach allows for a detailed analysis of nuclear structure by measuring the average pixel fluorescence intensity within each bin in the FITC and TRITC channels. For each bin, the max pixel fluorescence intensity was calculated across all the acquired images for both mutant and control groups for a single probe. The normalized intensity per bin was calculated for each cell line and each probe was determined using the minimum and maximum max pixel fluorescence intensities. This data was then used to create heatmaps and graphs on Prism10.

### RNA-seq analysis

Total RNAs were extracted from hiPSC-CMs and ventricular myocardial mouse tissues using RNeasy® Mini Kit Protocol (QiaGen, Cat#74104). Briefly, RNAs were extracted using Qiazol/chloroform solution, and purified with columns. The integrity of the RNA extracts was analyzed on an Agilent Bioanalyzer RNA chip and samples with RNA Integrity Number (RIN) read out of >8 were used. Strand-specific sequencing libraries were generated using the TruSeq stranded total RNA library preparation kit, including depletion of RNA (Illumina Inc.) and sequenced on NovaSeq6000 instrument (Illumina Inc.) to give 40 million reads of 100 bp, paired-end, by sample. The library preparation and sequencing were performed by the iGenSeq platform at Institut du Cerveau (Paris). The RNA-seq results were obtained using the following pipeline: reads were pre-processed using fastp module v0.20.0 and aligned on GRCh38 human or GRCMs39 mouse reference genome using star v2.7.5a. Pairs of reads aligned were annotated using gtf annotation files provided by ENSEMBL. Genome-aligned reads were indexed with samtools v1.13 and quantified with qualimap v2.2.2b. The number of reads uniquely mapping coding RNA was compiled in a counting matrix using featureCounts v2.0.1. All steps were submitted to quality control using mutliqc v1.13. Differential analysis was performed using the DESeq2 package on R using a threshold of p-adjusted (padj) value 0.05 and log2 fold change (log2FC) 0.5. Gene Ontology tables were obtained using Enrichr^58–60^, STRING v12.0 (https://string-db.org), and ggplot2 package on Rstudio.

### Protein extraction and immunoblotting

Total proteins were isolated from cardiac sections using an extraction buffer (Cell Signaling) containing protease and phosphatase inhibitors (Thermo Scientific™). The protein content of the samples was determined using the BCA Protein Assay Kit (Thermo Scientific™, #Cat23227). Protein extracts (50 µg) were analyzed by NuPAGE™ 4-12% Bis-Tris gels and transferred to nitrocellulose membranes (Invitrogen). After washing with Tris-buffered saline containing 1% Tween 20 (TBS-T), the membranes were blocked with 5% BSA in TBS-T for 1 h at RT and then incubated with the appropriate antibody at 4°C, O/N. Membranes were incubated with fluorescent-conjugated anti-mouse or anti-rabbit secondary antibodies (BioRad) for 1 h at RT. Antibody detection was imaged using the ChemiDoc Imaging System and ImageLab software (Bio-Rad). Quantification was performed using ImageLab software (Bio-Rad).

### Immunostaining of cardiac sections

Frozen heart tissue was cut into 10 μm thick cryosections, fixed (10 min, 4% PFA in 1X PBS at RT), and blocked (1% BSA, 5% goat serum, 0.2% Triton X-100 in 1X PBS) for 1 hour at RT. Sections were incubated with antibodies (1% BSA, 5% goat serum, 0.2% Triton X-100 in 1X PBS), O/N at 4°C, and then washed with 1X PBS. The sections were then incubated with the appropriate secondary antibodies for 1 h before being washed with PBS. Nuclei were counterstained with 0.005% Hoechst in 1X PBS for 5 min at RT. The slides were mounted with Fluoromount-G® (Invitrogen). Immunofluorescence microscopy was performed using an Invitrogen™ EVOS™ M5000 microscope, a fully inverted imaging system equipped with four-color fluorescence, transmitted light, and imaging capabilities. Images were acquired at ×20 objective with a pixel resolution of 0.44 µm/px. Images were processed using FIJI/ImageJ software.

### Electron microscopy

The cardiac sections were fixed in 2.5% glutaraldehyde diluted in PBS for 1 hour at room temperature. After washing in PBS, samples were post-fixed in 1% OsO4, dehydrated in a graded series of acetone, and embedded in epoxy resin. Ultrathin sections were cut at 90 nm, stained with uranyl acetate and lead citrate, examined with a transmission electron microscope (JEOL 1011), and photographed with an Erlangshen 1000 digital camera (GATAN) using Digital Micrograph software. The isolated cardiomyocytes were fixed in 2.5% glutaraldehyde in PBS for 3 h at RT. After PBS washes, 0.15 M of cacodylate was added and incubated at 40°C in a water bath. 3% of preheated agarose was added to the tube before solidification. The agarose-embedded cell pellet was cut into small pieces and incubated in a salt shaker into 0.15 M cacodylate, then 2% osmium tetroxide for 60 min at RT, light-protected. After distilled water washes, samples were dehydrated progressively in ethanol baths, then incubated two times with propylene oxide for 10 min at RT, and finally incubated in different epon resin baths before flat mold polymerization 48h at 50°C. Ultra-thin sections (80 nm) were obtained using ultramicrotome (UC7, Leica), and were contrasted with 2% uranyl acetate and citric acid lead. Primary observations were obtained using a transmission electron microscope with an acceleration voltage of 120 kV (JEOL 1400 flash, Japan) and a high-resolution digital camera (Xarosa, EM-SIS, Germany).

### Histological Analysis

Frozen hearts were cut into 8 μm thick sections and stained with Sirius red. Briefly, heart sections were fixed in 4% formaldehyde for 10 min, rinsed in EtOH 100%, dried for 20 min, and stained in 0.3% Sirius red solution for 1h. Then, sections were put in acetic acid 0.5% for 5 min twice, in EtOH 100% for 5 min three times, and finally in xylene for 10 min twice. Collagen fibrils were detected in red whereas cytoplasm remained yellow.

### RNA isolation and reverse-transcription qPCR

Total RNA was extracted from the mouse heart using RNeasy isolation kit (Qiagen) according to the manufacturer’s instructions. The adequacy and integrity of the extracted RNA were determined with the 2100 Bioanalyzer system (Agilent) according to the manufacturer’s instructions. cDNA was synthesized using the SuperScript III first-strand synthesis system according to the manufacturer’s instructions (Invitrogen). Real-time qPCR was performed with SYBR Green I Master mix (Roche) on the LightCycler 480 instrument (Roche). The primers utilized in this study are Col1A2-F CCGTGCTTCTCAGAACATCA, Col1A2-R GAGCAGCCATCGACTAGGAC, Col3a1-F TGTGGACATTGGCCCTGTTT, and Col3a2-R TGGTCACTTGCACTGGTTGA).

### Anti-Loxl2 treatment

Anti-Loxl2 (Abcam, Cat#ab0023) was administered by intraperitoneal (IP) injection to *Lmna^H^*^222^*^P/H^*^222^*^P^* mice at a dose of 5 mg/kg, three times a week for one month.

### Antibodies

The primary antibodies used for hiPSC-CMs immunostaining were mouse IgG1 anti-a-actinin (dilution 1:1000e, Sigma, Cat#A7732) and mouse IgG2b anti-cardiac Troponin I (cTnI) (dilution 1:1000e, Hytest, Cat#4T21). The secondary antibodies were goat anti-mouse IgG1 AlexaFluor 647 (dilution 1:1000, Invitrogen, Cat#A21240), goat anti-mouse IgG2b AlexaFluor 555 (dilution 1:1000e, Invitrogen, Cat#A21247). DNA was stained with DAPI (4’,6-diamidino-2-phenylindol, dilution 1:1000e).

The primary antibody used for Loxl2 immunostaining on mice cardiac sections was rabbit IgG anti-Loxl2 (dilution 1:100, Abcam, Cat#ab96300), anti-sarcomeric alpha-actin (Sigma, Cat#A2172). The secondary antibodies were goat anti-rabbit IgG AlexaFluor647 (Thermo Fisher Scientific, Cat#A21244) and goat anti-mouse IgM cross-absorbed AlexaFluor546 (Thermo Fisher Scientific, Cat#A21045). DNA was stained with 0.005% Hoechst.

The primary antibodies for the immunoblot were anti-Loxl2 (dilution 1:100, Abcam, Cat#ab96300) and anti-GAPDH (dilution 1:5000, SantaCruz - Cat#sc322331). The secondary antibodies used were goat anti-rabbit IgG StarBright Blue 520 (dilution 1:10,000, BioRad, Cat#12005870) and goat anti-mouse IgG StarBright Blue 700 (dilution 1:10,000, Cat#12004162).

### Statistics

The statistical significance of functional experiments on hiPSC-CMs was analyzed using a non-parametric Mann-Whitney statistical test. Cardiac echocardiography experiments in mice were analyzed using one-way ANOVA, with post-hoc comparisons as appropriate. Two-way ANOVA was used to analyze chromosome spatial organization, with post-hoc tests as needed. mRNA fibrosis marker expression was analyzed using an unpaired Student’s t-test. Loxl2 relative expression was quantified using an unpaired Student’s t-test. The effect of Loxl2 treatment on cardiac function, as measured by echocardiography, was analyzed using one-way ANOVA with Tukey’s multiple comparisons test. A significance level of p < 0.05 was used for all statistical tests. Data are presented as the mean ± SEM, and sample sizes are indicated in the figure legends. Statistical analyses were performed using Prism software (GraphPad, version 9).

## ACKNOWLEDGEMENTS

This work was supported by the Association Française contre les Myopathies, Région Occitanie, and ANR Musage. We thank the genomic facility iGenSeq at the Brain Institute, Paris, for the ATAC-seq and RNA-seq libraries sequencing. We thank Dr. Ben Yaou (Institute of Myology Paris) and Dr. Eschenhagen (University Medical Center Hamburg-Eppendorf) for providing the hiPSCs carrying the *LMNA* p.H222P mutation. We Thank Zoheir Guesmia from MyoImage, Myology Institute, Paris, for his help with image analysis and pipeline development.

## AUTHOR CONTRIBUTIONS

Conceptualization, AM and ACM; investigation, MK, CP, FG; cardiomyocytes isolation, AF; cardiac function assessment, NM and AM; hiPSC, MK, AC and ACM; images analysis and pipeline development, ZG; electron microscopy, GB; writing - original draft, AM; writing - review and editing, MK and ACM; funding acquisition, AM and ACM; supervision, AM and ACM.

---

**Figure S1:**
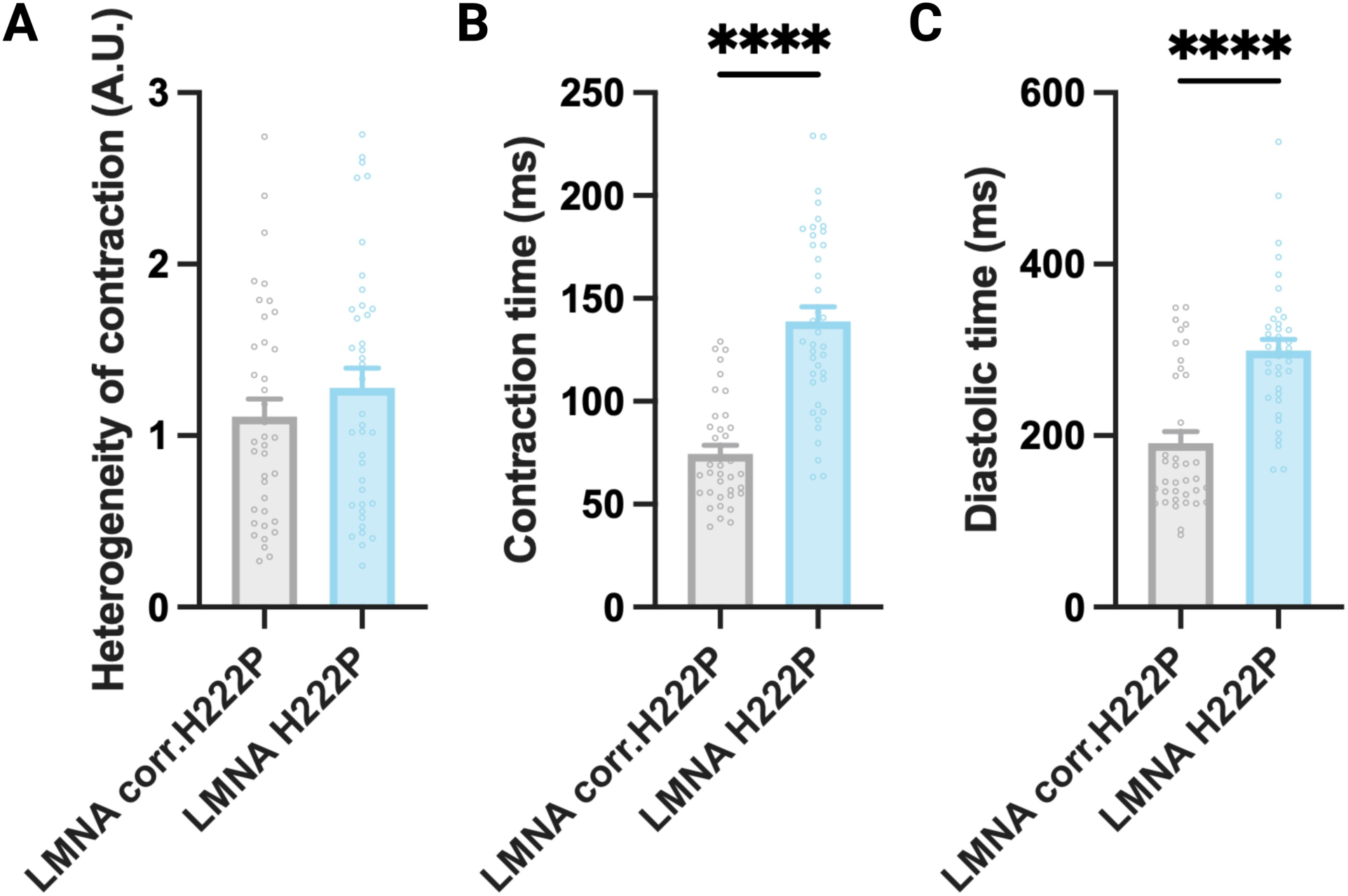
Measure of the contractile properties under pacing in hiPSC-CMs. Measure of the **(A)** heterogeneity of contraction, **(B)** contraction time (*** p <0.0001), and **(C)** diastolic time (*** p <0.0001). (Mann-Whitney statistical test, mean ± SEM; nH222P= 3, ncorr.H222P= 3).

**Figure S2:**
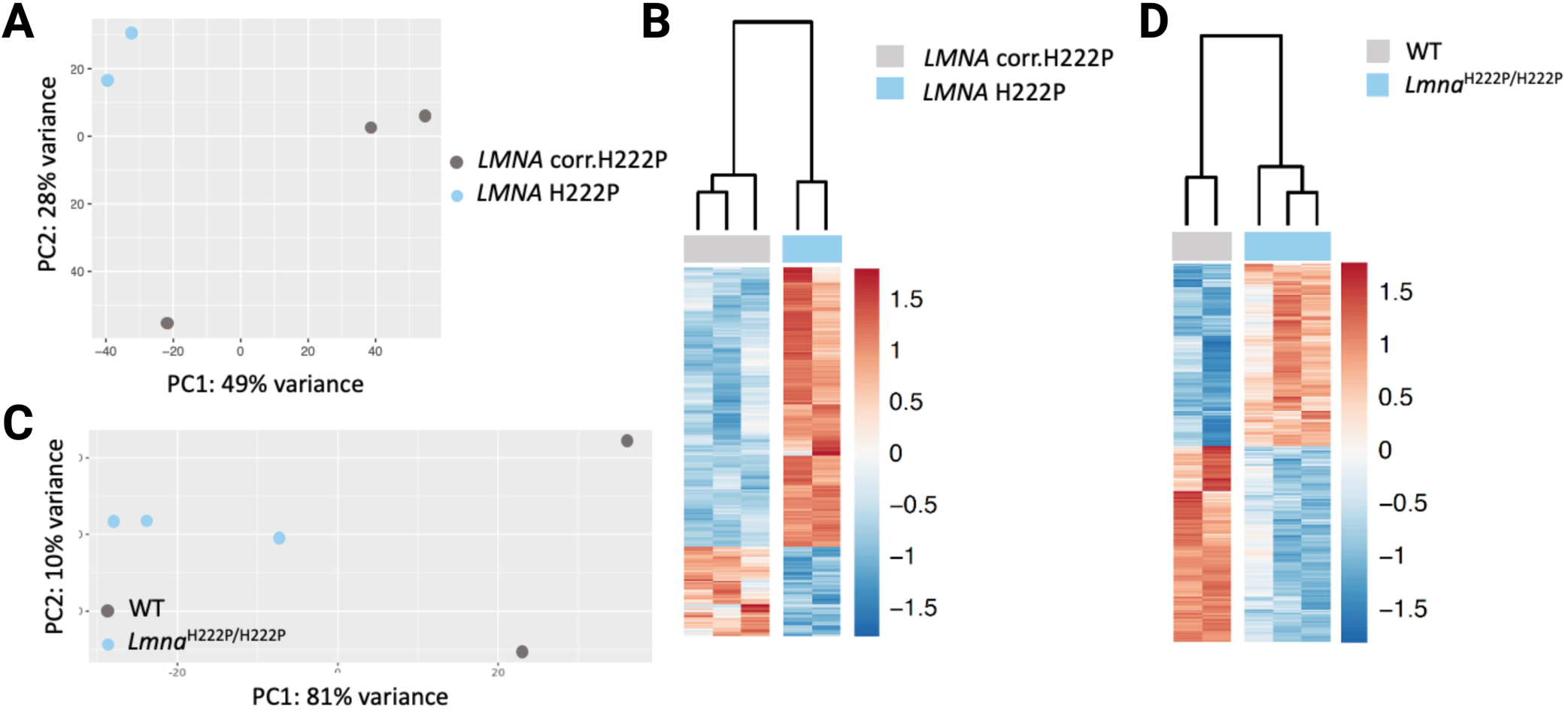
Transcriptomics analysis in both *in vitro* and *in vivo*. **(A,C)** PCA of **(A)** hiPSC-CMs and **(B)** mice. **(B,D)** Heatmaps representing DEGs identified in **(B)** hiPSC-CMs and **(C)** mice. (nH222P= 2, ncorr.H222P= 3, n*Lmna*= 3, nWT= 2; threshold, padj 0.05, log2FC 0.5).

## Notes

### Competing Interest Statement

The authors have declared no competing interest.

